# A Rapid and High-Sensitivity Non-radioactive Screening Assay for Insulin Receptor Ligands

**DOI:** 10.1101/2025.03.20.644157

**Authors:** Jana Staňurová, Kateřina Čermáková, David Just, Ullrich Jahn, Benjamin Fabre, Erik Rozkovec, Lenka Žáková, Pavel Šácha, Jiří Jiráček, Jan Konvalinka

## Abstract

Insulin is a key hormone in glucose homeostasis. Its lack causes severe health complications and has to be compensated by regular administration of insulin. Despite intense long-lasting research, a more stable and efficient substitute has yet to be discovered to alleviate patients’ issues. Here we report the development of a new assay for screening potential insulin receptor ligands based on the DNA-linked inhibitor antibody assay (DIANA). Our assay meets the need for a fast, sensitive, non-radioactive method as an alternative to the commonly used radioligand receptor binding assay.

## Introduction

The key function of insulin is to regulate the uptake of glucose from the blood ^[1]^. Upon detection of elevated glucose levels, secretion of insulin from the pancreas is quickly initiated. Signaling triggered by binding of the hormone to its receptor ^[2]^ leads to the relocation of glucose transporter to the plasmatic membrane of target cells, i.e. muscle and adipose cells, facilitating the uptake of glucose by these cells. The full function of insulin-insulin receptor pathway is crucial for glucose homeostasis and metabolism.

Diabetes mellitus type I is caused by the autoimmune destruction of pancreatic β-cells ^[3]^. As a result, insulin production is lost and glucose homeostasis is disrupted, leading to characteristic acute symptoms such as polyuria, polydipsia, polyphagia, unexplained weight loss, fatigue, and blurred vision. If untreated, the disease may rapidly progress to life-threatening diabetic ketoacidosis, which can result in nausea, vomiting, abdominal pain, deep labored breathing, and eventually coma and death ^[4]^. Administration of insulin remains the only life-saving treatment for patients with type I diabetes, with subcutaneous delivery compensating for the loss of endogenous insulin production, so that glucose levels in blood remain under control, preferably at the physiological level of healthy individuals ^[5]^. However, the need for daily s.c. medication deteriorates the quality of the patients’ lives substantially. An insulin substitute that is more stable, cheaper and potentially orally available would significantly alleviate this burden ^[6]^. We therefore aimed to establish a new method capable of screening a large number of compounds simultaneously for their ability to bind to the insulin receptor. This method may prove useful for exploring available compound libraries in search for novel insulin receptor ligands.

We employed DNA-linked inhibitor antibody assay (DIANA), a technique developed in our laboratory, to perform compound screening (Figure 1) ^[8]^. DIANA is a sandwich-based assay that relies on the interaction of a selected target protein with its ligand. The ligand is usually a small molecule known to interact with the protein of interest. It is conjugated to a DNA oligonucleotide forming the DIANA detection probe (Figure 2). Quantification is executed via RT-qPCR. In the competitive setting of the method, the probe is mixed with tested compounds (ligands), which leads to a decrease in the amount of bound probe. The K_d_ value is then determined based on the compound concentration in a single well.

**Figure 1:**
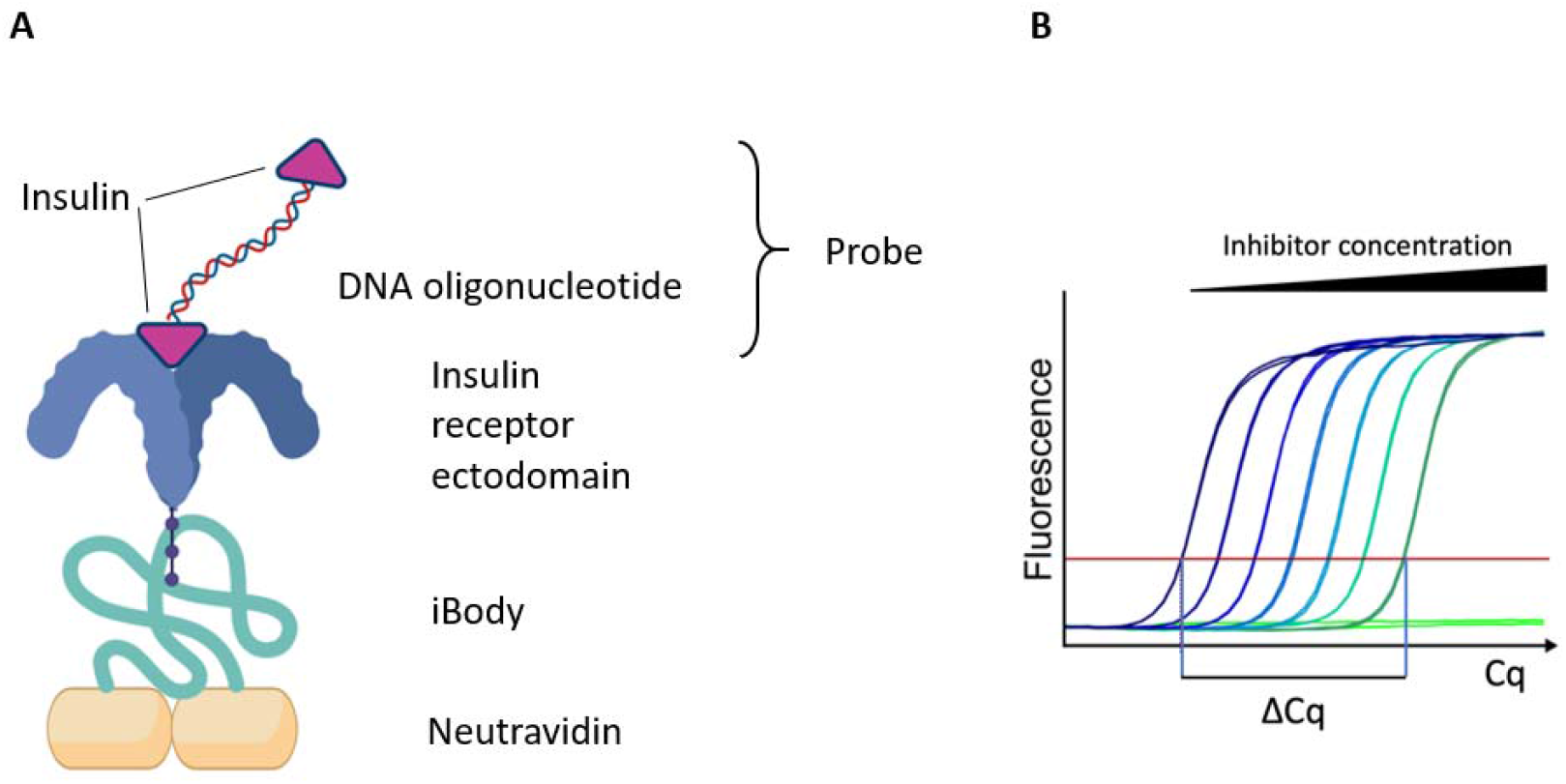
Schematic representation of DIANA. **A**. iBody, a HPMA polymer decorated with tris-nitrilotriacetic acid for the binding of a His-tag, and biotin ^[7]^ facilitates a stable immobilization of the insulin receptor ectodomain to a neutravidin-coated surface. Detection probe comprising two insulin moieties attached to one DNA oligonucleotide binds to the insulin receptor via its insulin molecule. **B**. The amount of detection probe bound to the IR is measured by qPCR. Higher concentrations of an inhibitor lead to a more pronounced displacement of the detection probe resulting in a higher measured Cq (cycle of quantification). ΔCq, termed as dynamic range of the assay or assay window, represents the difference between an uninhibited reaction (left) and a fully inhibited reaction (right). Created in BioRender. Stanurova, J. (2025) https://BioRender.com/pkj691y

**Figure 2:**
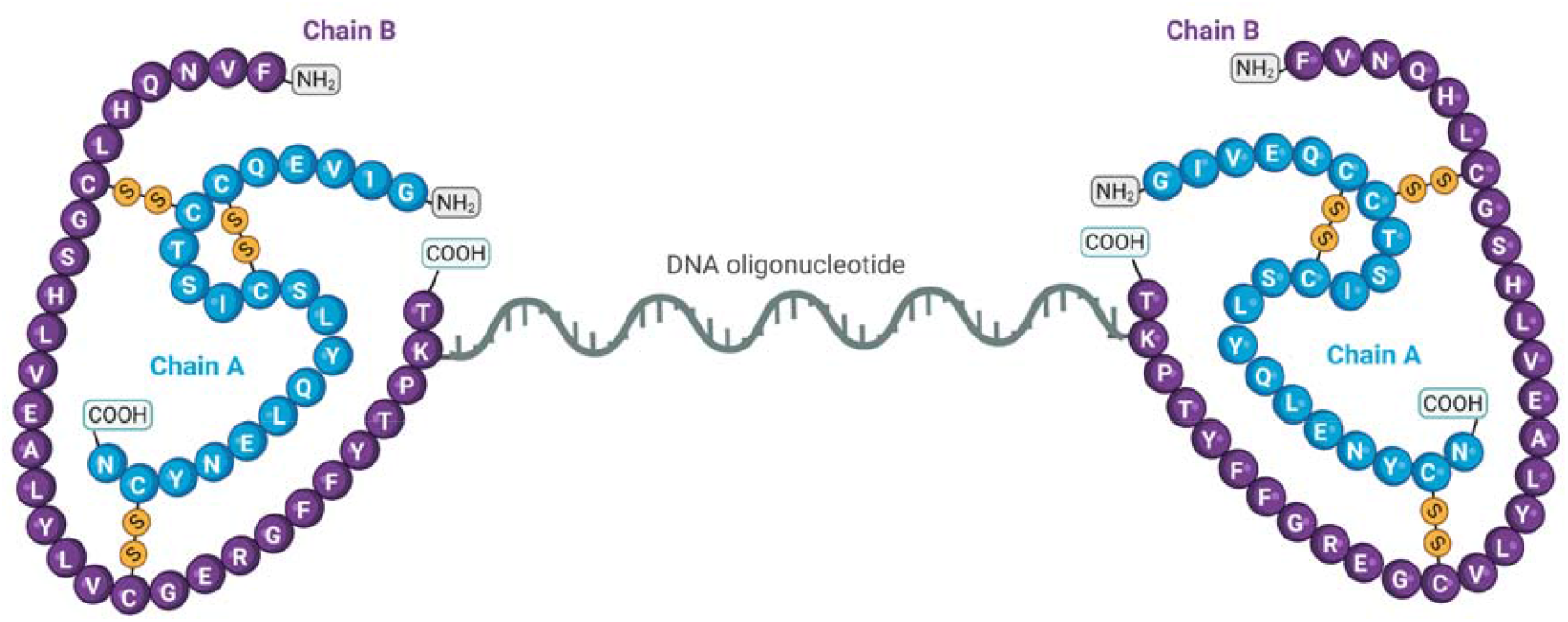
Schematic representation of the insulin detection probe. Two insulin molecules are attached to each end of the DNA oligonucleotide. Created in BioRender. Stanurova, J. (2025) https://BioRender.com/mpfypma

As opposed to radioimmunoassay (RIA) ^[9], [10]^ utilized to measure insulin concentration in samples routinely, and radioligand receptor binding assays ^[11]^ used to determine ligand affinity for the insulin receptor, there is no need for radioactive labeling of insulin in DIANA. Also, the method is not cell-based, thus minimizing biological variability in the results. So far, our laboratory successfully developed DIANA for screening of glutamate carboxypeptidase II (GCPII)^[8]^, carbonic anhydrase IX (CAIX)^[12]^, neuraminidase^[13]^, fibroblast activation protein (FAP)^[14]^ and cluster of differentiation 73 (CD73)^[15]^ inhibitors.

## Methods

### Synthesis of the B29-azidolysine-insulin

The precursor for the synthesis of the DIANA probe, B29-azidolysine-insulin, was prepared by the trypsin-catalyzed semisynthesis from [*des*(B23-B30)]-insulin (DOI) and GFFYTPK(εN_3_)T peptide (where εN_3_ is azidolysine) according to Žáková et al. GFFYTPK(εN_3_)T peptide was synthesized by the solid phase synthesis according to Viková et al. ^[17]^.

### DIANA probe preparation and characterization

DIANA probe is a conjugate of a ligand, B29-azidolysine-insulin in our case, with a DNA oligonucleotide. The modification of a ligand is a key step in the preparation of DIANA probe. Modification of B29 position in insulin with an azido group allows conjugation with an alkyne-modified DNA oligonucleotide by copper(I)-catalyzed azide-alkyne cycloaddition (CuAAC). Insulin derivative was added in 3-fold excess (100 μM) to a 78-nucleotide-long DNA oligonucleotide (33 μM) and the mixture with 5 mM aminoguanidine, 5 mM BTTP, 2.5 mM CuSO_4_ and 20 mM natrium ascorbate was incubated at 37 °C for three hours. The product, a dimer of two insulin molecules linked at their B29 positions via a DNA oligonucleotide, was purified using a filter column (Amicon, Merck), followed by LC-MS analysis. The probe was characterized by determining linear range and K_d_ value for the immobilized insulin receptor. Linear range was assessed as a serial dilution of insulin receptor (0.1-1500 pg/well) detected with 100 pM DIANA probe (Figure S1A). K_d_ value was measured as 1500 pg/well of insulin receptor detected with probe dilution series (40 fM – 20 μM) and calculated according to Navrátil et al. ^[8]^ (Figure S1B). The affinity of the probe to insulin receptor, K_d_ = 0.6 ± 0.1 nM, is similar to the affinity of insulin alone with K_d_ = 1.1 ± 0.2 nM (Figure S1C). Binding capacity of insulin was not impaired post attachment to the DNA oligonucleotide.

The determination of K_d_ is a fundamental assessment in the evaluation of interactions between two molecules in binding assays ^[18]^. In the context of DIANA, it serves as a crucial parameter for determining the K_d_ values of inhibitors in a single experiment ^[8]^. As a basic parameter, K_d_ is influenced by factors such as assay setup, assay type, buffer composition, pH, and assay sensitivity, meaning its value can vary across different methods ^[18], [19]^.

### DIANA optimization and validation for screening

To ensure successful measurement of ligands, the DIANA experimental conditions were optimized. Factors such as blocker type, detergent type, their concentrations, and incubation times all impact key experiment parameters like dynamic range, reproducibility, accuracy, and equilibrium state.

For optimization, we tested various buffer compositions with different concentrations of casein and detergents (Tween-20 and Pluronic F-127). The conditions for ligand screening were chosen based on the highest signal-to-background ratio (ΔCq, Figure 1B) and assay sensitivity. After selecting the appropriate buffer, we proceeded with optimizing the concentrations of protein and probe. The optimal IR/probe concentration ratio was determined by the highest ΔCq, which is the difference between the Cq value at a high concentration of insulin and the Cq value of wells without insulin. The final optimal conditions for competitive DIANA were 0.01% Tween-20, 700 pg IR per well, 50 pM bivalent DIANA detection probe, and one-hour incubation with inhibitors.

Given that most compounds in libraries are dissolved in DMSO, we also examined the impact of DMSO on our assay. Since many targets tested in HTS assays are sensitive to higher DMSO concentrations, we focused on conditions, in which most compounds would remain soluble to avoid false-negative results from precipitates. We selected the highest DMSO concentration that did not significantly reduce the assay’s dynamic range or affect K_d_ values compared to DMSO-free experiments. Therefore, a 10 % DMSO concentration was chosen for inhibitor screening, including HTS (Figure S2).

### DIANA conditions

DIANA in a competitive setting was developed to determine the compounds’ binding potency towards the insulin receptor. The assay was performed in 96-well plates with a working volume of 5 μl per well, except for casein. Neutravidin was immobilized on polypropylene well plate surface (5 μl, 12 ng/μl) in TBS (20 mM Tris-HCl, 150 mM NaCl, pH 7.5) and incubated for 1 hour. Blocking by 100 μl 1.1 % casein solution in TBS overnight followed. Next, iBody decorated with TrisNTA and biotin ^[7]^ was applied to the plate (5 μl, 30 nM) in TBST’ (20 mM Tris-HCl, 150 mM NaCl, 0.1 % Tween-20, pH 7.5) and incubated for 1 hour. In the following step, insulin receptor (IR-A ectodomain with His-tag, SinoBiological, 11081-H08H) was added to the plate (5 μl, 700 pg/well) in TBST’ and left for 1 hour. Afterwards, detection probe was applied to the plate with or without tested compounds (5 μl, 50 pM) in HEPEST’C (100 mM HEPES, 400 mM NaCl, 0.1 % Tween-20, pH 7.5) complemented with 0.0055 % casein. All incubations were carried out at room temperature in the dark. After each step, the plate was washed by automated plate washer (BlueWasher, BlueCatBio) with TBST (20 mM Tris-HCl, 150 mM NaCl, 0.05 % Tween-20, pH 7.5). Finally, master mix (5 μl, LightCycler^®^ 480 SYBR Green I Master, Roche) was added. RT-qPCR ensued, quantification analysis was performed, and K_d_ values were calculated as described in Navrátil et al. ^[8]^.

### Synthesis of ligands

The full-length insulin analogs [*cyclo*(Nva(δN_3_)B26, PrgB29]-insulin (**LZ-90**), [*cyclo*(*D*-Nva(δN_3_)B26, *D*-PrgB29]-insulin (**LZ-135**), [*cyclo*(Nva(δN_3_)B26, *D*-PrgB29]-insulin (**LZ-137**), and [*cyclo*(*D*-Nva(δN_3_)B26, PrgB29]-insulin (**LZ-136**) contain side chains at positions B26 and B29 connected by a triazole-containing bridge and were prepared as described by Viková et al. ^[17]^.

[GlyB31, B31-carboxamide]-insulin (**LZ-124**), featuring a B-chain C-terminus extended by glycine carboxamide, was prepared as described by Páníková et al. ^[20]^.

The proline derivative-containing analogs [*des*(B28–B30), *D*-ProB26, B26-carboxamide]-insulin (**LZ-59**), [*des*(B28–B30), *D*-ProB26, B27-carboxamide]-insulin (**LZ-60**), [*D*-ProB26]-insulin (**LZ-61**), [4-(*R, trans*)-hydroxy-ProB26, B30-carboxamide]-insulin (**LZ-122**), and [4-(*R, trans*)-hydroxy-ProB26, 4-(*R, trans*)-hydroxy-ProB28, B30-carboxamide]-insulin (**LZ-123**) were synthesized as described by JiráČek et al. ^[21]^.

The next series of ligands includes one full-length insulin analog (**LZ-113**) and two des(B27– B30)-shortened analogs (**LZ-120** and **LZ-128**), all featuring sterically hindered, non-canonical amino acids at the B26 position of insulin. The structures of these amino acid building blocks are shown in Figure S3, and the synthesis of their precursors as well as the corresponding insulin analogs is described in the Supporting Information (SI).

The cyclic 20-amino-acid-long insulin mimetics **JZ-2-056SS** and **ML-074** (Figure 5) contain either a disulfide or a dicarba bridge at positions 11 and 18, respectively, and were prepared as described by Lubos et al. ^[22]^.

The insulin mimetics **MGH31-2-1** and **MGH-131-1** contain two different peptide sequences attached via their C-termini to a central adamantane or trimesic acid scaffold, respectively (Figure 6). **MGH31-2-1** was prepared according to Hajduch et al. ^[23]^ and **MGH-131-1** as described in Selicharová et al. ^[24]^.

### Determination of binding affinity to the insulin receptor (IR) by radioligand assay

The radioligand binding assay with the human insulin receptor (IR-A isoform) in IM-9 cells (ATCC) was conducted following the method outlined by Morcavallo et al. ^[25]^. This assay is based on the competition between unlabeled insulin and radiolabeled insulin for binding to the insulin receptor in living cells. Initially, the standard binding curve for recombinant human insulin to the human insulin receptor in IM-9 cells was determined. In brief, 2.0 × 10^6^/ml IM-9 cells were incubated with increasing concentrations of human insulin and a constant concentration of human [^125^I]monoiodotyrosyl-A14-insulin (prepared according to Asai *et al*. 10.1007/s00216-021-03423-3) for 2.5 hours at 15 °C in HEPES binding buffer (100 mM HEPES, 100 mM NaCl, 5 mM KCl, 1.3 mM MgSO_4_, 1 mM EDTA, 10 mM glucose, 15 mM NaOAc, 1 % BSA, pH 7.6), with a total volume of 500 µl. After incubation, 2 x 200 µl samples were centrifuged at 13,000 x g for 10 minutes. Radioactive pellets were counted using a Wizard 1470 Automatic γ Counter (PerkinElmer Life Sciences). The binding data were analyzed using GraphPad Prism 8 with a non-linear regression method and a one-site fitting program, accounting for potential depletion of the free ligand, which allowed the determination of the dissociation constant for the unlabeled ligand. The dissociation constant of human [^125^I]monoiodotyrosyl-A14-insulin was set at 0.3 nM.

### HTS screening

Screening was performed to determine K_d_ values using the DIANA method, as previously described Tykvart et al. ^[12]^, under the same conditions as noted in DIANA conditions with minor modifications: blocker solution (1.1 % casein in TBS) was added to neutravidin at 20 µl per well; in the probe step, a mixture of 10 µM tested compounds and the bivalent DIANA detection probe was added to the insulin receptor (5 µl per well) in HEPES buffer containing 0.05% Tween-20 and 0.0055% casein. The probe-inhibitor mixture was incubated for one hour. After a final wash, qPCR mix was added for probe detection.

HTS was performed in an automated HEPA-filtered workcell. The instruments used included Agilent Bravo Automated Liquid Handling Platform for neutravidin coating, and iBody, insulin receptor and probe addition; Formulatrix Mantis^®^ Microfluidic Liquid Dispenser for casein and qPCR master mix application; Beckman Coulter Echo 650 Acoustic Liquid Handler for transferring library compounds; BlueCatBio BlueWasher for plate washing; and Roche LightCycler 480 II for RT-qPCR runs.

The K_d_ value of a ligand can be determined from a single-well experiment ^[8]^. In case of insulin receptor, we prepared dilution series of each promising ligand and calculated the K_d_ values from a point within the dynamic range of the assay and evaluated them statistically as a mean of assessed K_d_ ± SD.

### Chemical libraries

Screening of two libraries in an HTS format was performed. The OTAVA Alpha Helix Peptidomimetic Library contains 1298 compounds (otavachemicals.com) and it is of significant value in drug discovery projects focused on protein-protein interactions. The peptidomimetic compounds were designed by modeling based on published results considering molecular descriptors such as LogP, molecular weight, number of hydrogen donors and acceptors, amount of rotatable bonds, quantity of rings, and molecular polar surface area.

The IOCB library is a collection of compounds synthesized at the Institute of Organic Chemistry and Biochemistry of the Czech Academy of Sciences in an effort of developing successful molecules in the field of medicinal chemistry. It contains 10560 substances of diverse chemical structures, such as antiviral drugs, steroids, natural products, nucleotide and nucleoside analogs etc.

## Results and discussion

We performed high-throughput screening of two libraries – the IOCB library comprising 10560 compounds in a pooled format (Figure 3) and the OTAVA Alpha Helix Peptidomimetic Library containing 1298 compounds. The assay window of the HTS experiments was 8.2 cycles and 7.3 cycles, respectively. The latter library did not yield any hits worth pursuing, since their K_d_ values were over 100 µM. The former library yielded several hits, all of which were insulin analogs (**LZ-90 – LZ-128**) or insulin mimetics (**JZ-2-056SS, ML074, MGH31-2-1** and **MGH131-1**) (Table 1) previously prepared by us. This is an expected outcome, since both libraries are rather small. Especially the IOCB library was established for different targets than the IR, mostly focusing on catalytic sites of various enzymes. It is a collection of various compounds synthesized at the IOCB and contains many nucleoside derivatives, which reflects the history of IOCB in developing inhibitors of this nature.

**Table 1.** Comparison of K_(d)i_ values determined by radioligand binding assay with IR-A in cells and DIANA. Compounds marked in the table by * represent those identified by HTS screening. For derivatives indicated by ^#^, their K_d_ measured by DIANA is notably lower than that determined by radioligand binding assay with IR-A in cells. Conversely, compounds specified by ^&^ are those with K_d_ values assessed higher in DIANA than in IR-A.

| Code | $K_d$ (nM ± S.D.)<br>for radioligand<br>binding assay<br>(n) | $K_d$ (nM ± S.D.)<br>From DIANA | $K_d$ DIANA/$K_d$ binding<br>assay with IR-A in<br>cells |
| --- | --- | --- | --- |
| Human insulin | $0.30 \pm 0.01$ (4) | $0.91 \pm 0.09$ | 3.0 |
| LZ-90 | $0.19 \pm 0.02$ (3) | $0.77 \pm 0.22$ | 4.1 |
| LZ-135 | $1.07 \pm 0.32$ (3) | $4.29 \pm 0.11$ | 4.0 |
| LZ-137 & | $0.25 \pm 0.09$ (3) | $1.95 \pm 0.24$ | 7.8 |
| LZ-136 & | $1.8 \pm 0.04$ (3) | $10.6 \pm 0.2$ | 5.9 |
| LZ-124 | $0.53 \pm 0.11$ (3) | $2.17 \pm 0.62$ | 4.1 |
| LZ-59 | $0.27 \pm 0.07$ (4) | $0.74 \pm 0.26$ | 2.7 |
| LZ-60 | $3.04 \pm 0.22$ (3) | $12.68 \pm 1.68$ | 4.2 |
| LZ-61 # | $104 \pm 61$ (3) | $93 \pm 23$ | 0.9 |
| LZ-122 & | $3.76 \pm 1.28$ (3) | $25.75 \pm 1.41$ | 6.8 |
| LZ-123 | $1.06 \pm 0.36$ (3) | $5.52 \pm 0.32$ | 5.2 |
| LZ-113 | $3.65 \pm 0.48$ (3) | $16.43 \pm 2.76$ | 4.5 |
| LZ-120 | $3.82 \pm 1.18$ (3) | $10.17 \pm 0.59$ | 2.7 |
| LZ-128 | $1.02 \pm 0.40$ (3) | $3.79 \pm 0.24$ | 3.7 |
| JZ-2-056SS * | $3.9 \pm 1.0$ (3) | $8.1 \pm 1.3$ | 2.1 |
| ML074 * | $49 \pm 6$ (3) | $102 \pm 16$ | 2.1 |
| MGH31-2-1 *# | $395 \pm 27$ (3) | $124 \pm 34$ | 0.31 |
| MGH131-1 *# | $1260 \pm 244$ (3) | $73 \pm 8$ | 0.06 |

**Figure 3:**
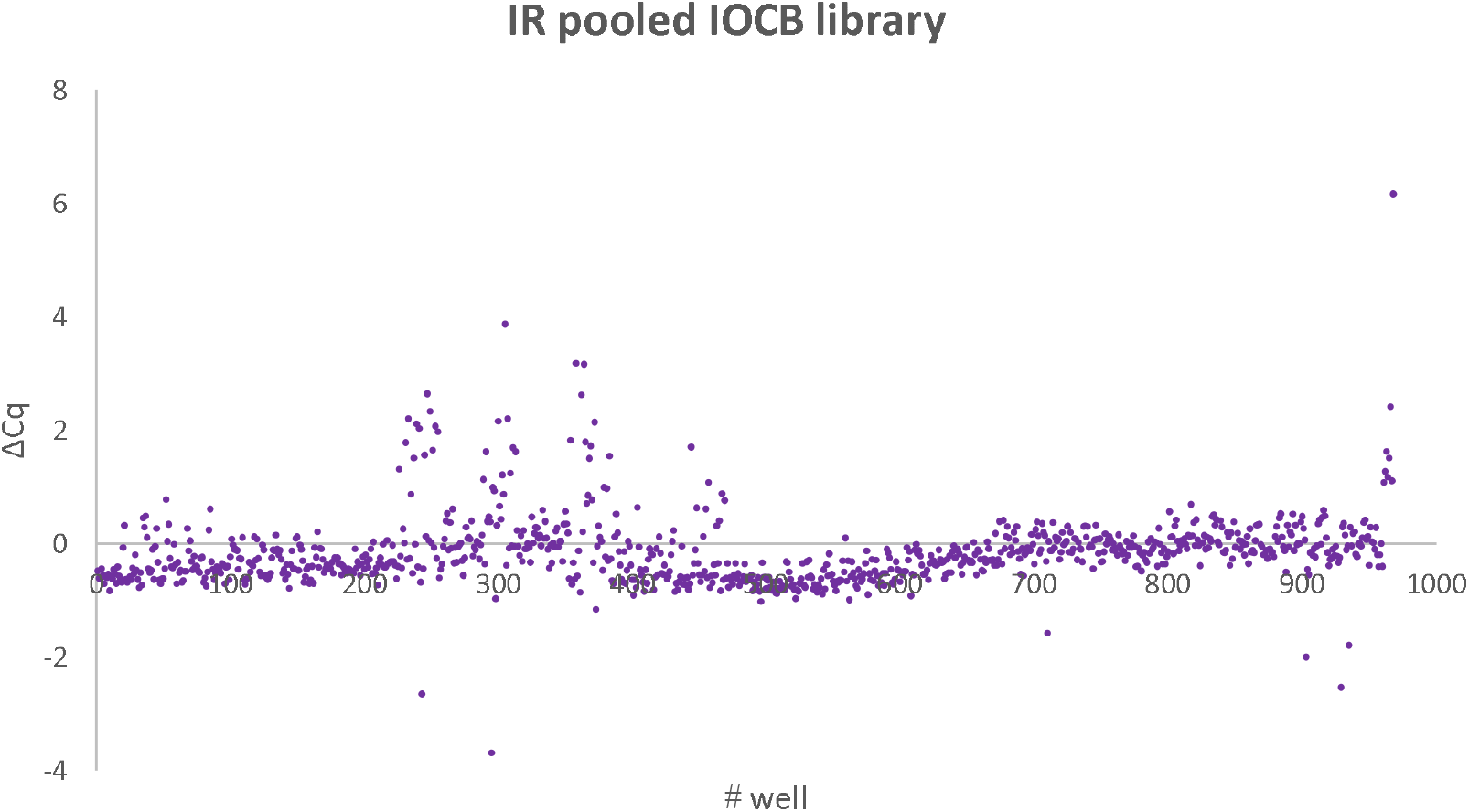
Results of the pooled IOCB library screening against IR. On the x axis, wells containing 11 compounds are plotted, the y axis represents ΔC_q_ values (the difference between the mean C_q_ of wells without any compound and the C_q_ value for each specific well). Wells with ΔC_q_ > 1 were evaluated as hits. Values of ΔC_q_ < −1.0 were excluded as interferences.

Nonetheless, insulin DIANA is a valuable tool for screening. It is a fast, sensitive method with very good reproducibility, non-radioactive and easily scalable to a HTS format. Moreover, cells are not necessary to perform the measurement, which reduces variability in the results. Another noteworthy merit of the method is the low consumption of material making the assay cost-effective in addition. With DIANA, we are able to reproduce results from IR-A to a good extent (Figure 4). In general, with a few exceptions, the K_d_ values determined by DIANA were higher than K_d_ values assessed by radioligand binding assay with IR-A in cells. For the insulin analogs (**LZ-90** – **LZ-128**, Table 1), the affinities determined by DIANA are fairly consistently about 4.4-fold higher (4.35 ± 0.35, n = 13) than the values determined using the cell-based IR-A binding assay. Only analog **LZ-61** clearly deviates from this trend, showing similar affinity values by both methods. However, **LZ-61** is a markedly weak binder compared to the other analogs. This suggests that weak binding to the insulin receptor may account for the “anomalous” behavior of K_d_ values determined by the two methods: the weaker the analog, the greater the discrepancy observed. It further appears that the DIANA assay has a tendency to overestimate the affinity (i.e., underestimate K_d_ values) of very weak binders.

**Figure 4:**
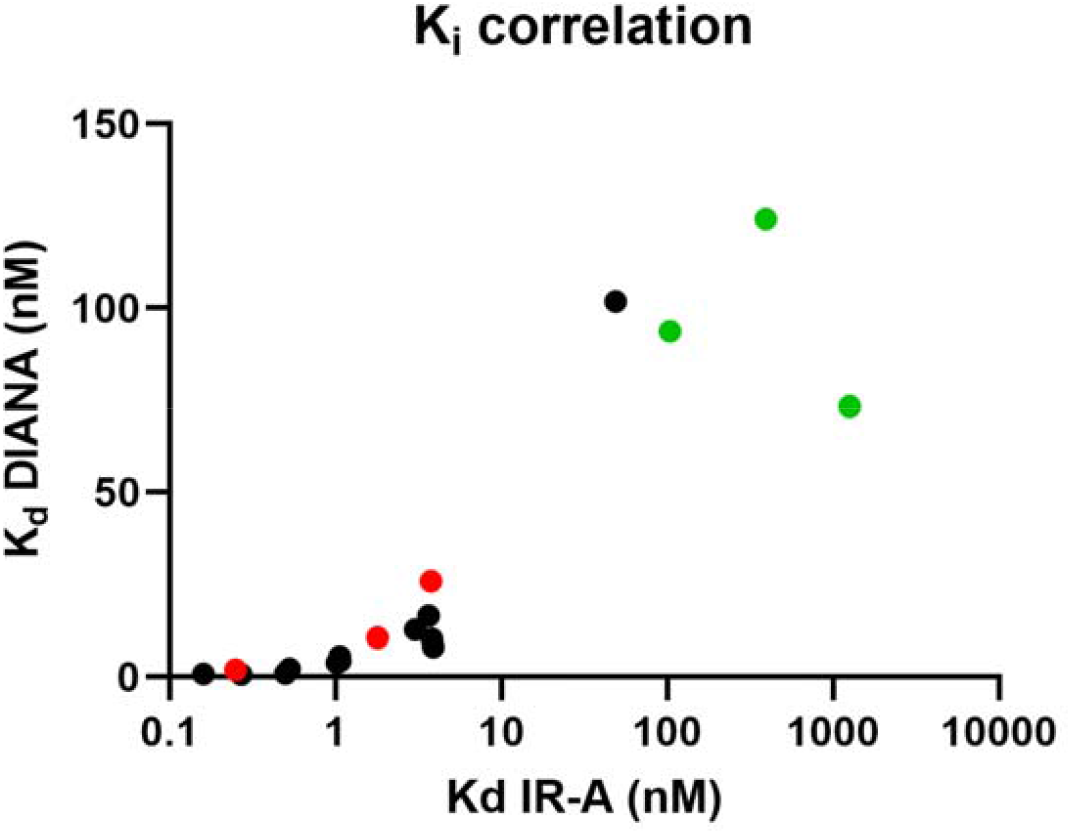
Correlation of K_d_ values for compounds listed in Table 1 measured by radioligand binding assay with IR-A in cells (x axis) and DIANA (y axis). Substances represented in green correspond with those marked by ^#^ in Table 1, that is those with lower K_d_ values determined by DIANA than by radioligand binding assay. Red dots denote derivatives with an opposite result, compounds specified by ^&^ in the table above. Molecules with a very good correlation of K_d_ values assessed by both methods are indicated in black. R^2^ = 0.8007, semilog nonlinear fit (x axis log, y axis linear).

**JZ-2-056SS** and **ML074** are 18-residue cyclic peptides ^[22]^, whose structures and sequences (see Figure 5) bear no resemblance to insulin. In contrast to insulin, which binds to both site 1 and site 2 of the insulin receptor, these peptides have been reported in the literature to bind exclusively to site 2 ^[22]^. Their affinities measured by DIANA are about 2-fold higher than the affinities determined by the radioligand binding assay. The reason why non-insulin-derived compounds exhibit a different affinity ratio compared to insulin analogs is not obvious, but it likely relates to their distinct mechanism of receptor binding.

**Figure 5.**
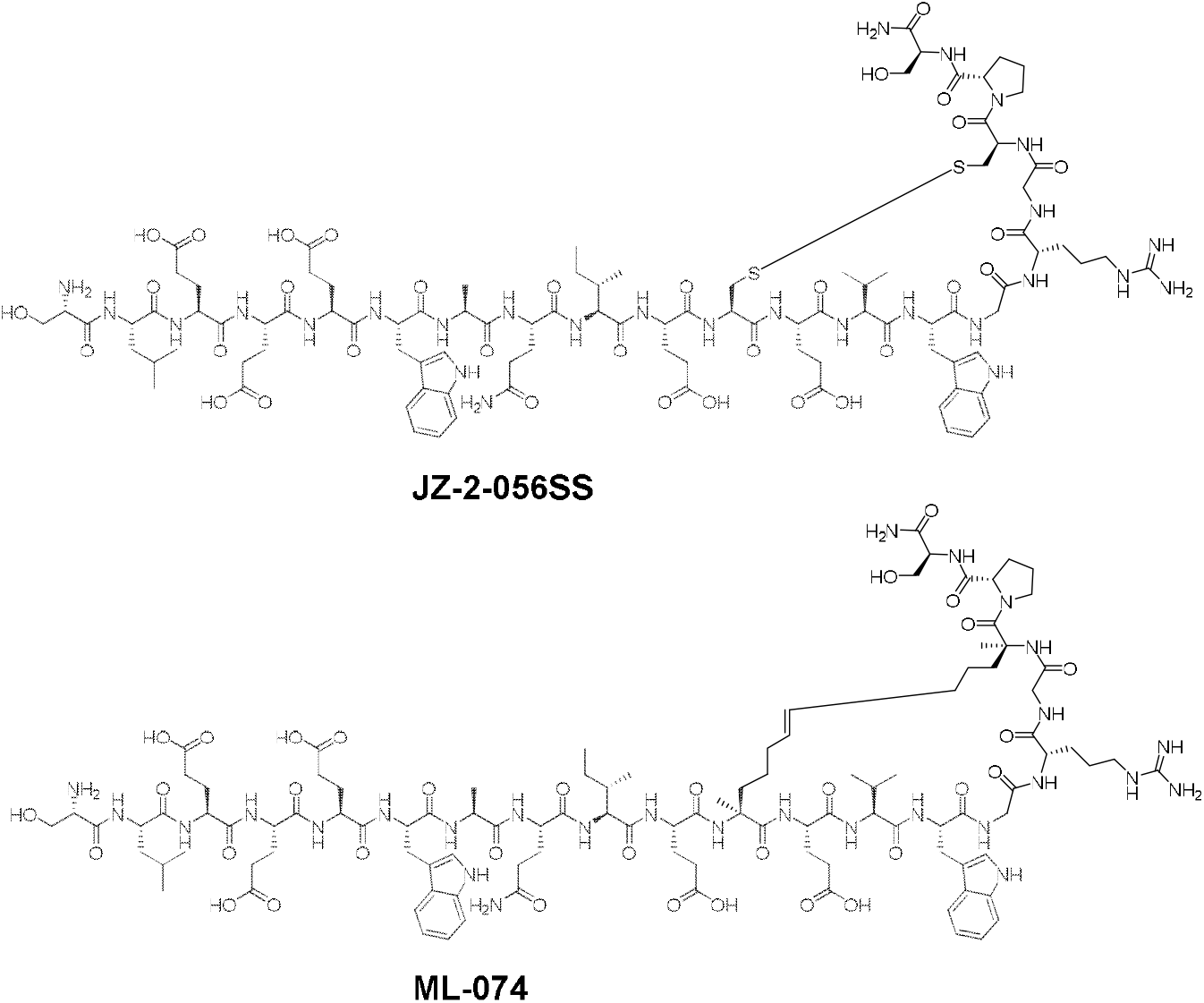
Structures of insulin mimetics JZ-2-056SS and ML074 ^[22]^.

**MGH31-2-1** and **MGH131-1** are insulin mimetics ^[23], [24]^, in which peptide sequences specific for both site 1 and site 2 of the IR are covalently attached by their C-termini to a central non-peptidic scaffold (structures provided in Figure 6). Binding of these compounds to the receptor is significantly weaker than that of the other tested compounds, including **LZ-61**. Only for these two compounds, **MGH31-2-1** and **MGH131-1**, the assessed K_d_ values were notably lower in DIANA than in radioligand binding assay. We found that **MGH31-2-1** interferes with the qPCR, most likely by binding to the probe and hindering the elongation phase. This results in an artificially low K_d_ value, so we conclude the K_d_ measurement performed by radioligand binding assay on the cell is the more reliable one. The compound does not seem to act as an intercalating agent, since no interference occurred in qPCR with the oligonucleotide used in the click reaction. However, due to the size and structure of the ligand, it is conceivable that it binds to the protein part of the probe preventing the correct formation of the DNA duplex. The latter ligand, **MGH131-1**, was tested in the same settings as the former one. It did not interfere with the qPCR of the oligonucleotide alone. Interestingly, we did not observe any effect on qPCR with the probe either, so the interference must take place within the DIANA sandwich, yet we do not know the exact mechanism. A similar mechanism may be behind the marked differences in K_d_ values measured by DIANA for other weak binders like **LZ-61**.

**Figure 6:**
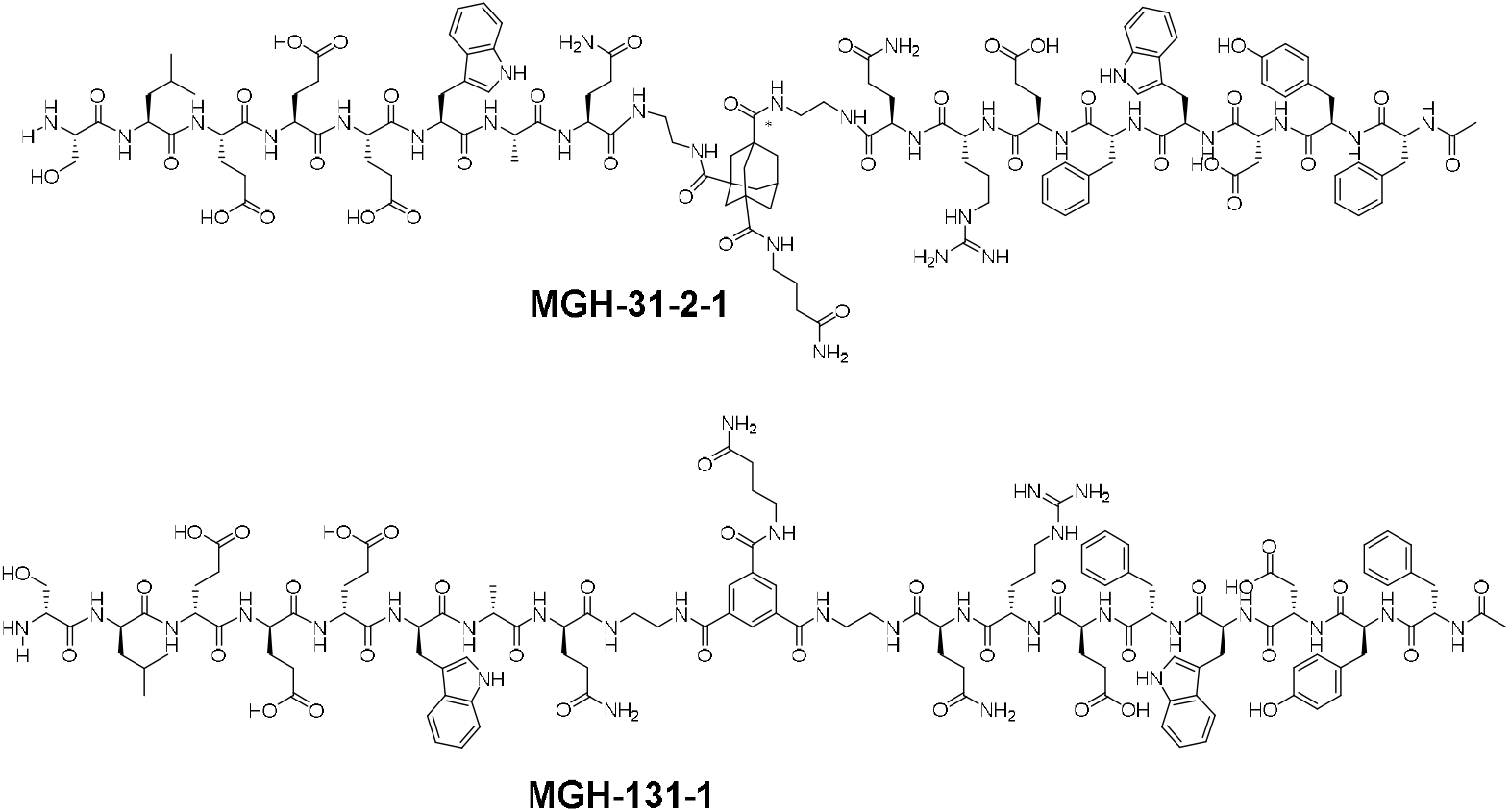
Structures of insulin mimetics MGH31-2-1 ^[23]^ and MGH131-1 ^[24]^.

The difference in binding affinities determined by the two methods may also be caused partly by different conditions and partly by the setups themselves. In radioligand cell-based assay, buffer is added first, then the tested compounds, then does the [^125^I]monoiodotyrosyl-A14-insulin follow and at the end, the cells are added. This way, the insulin receptor is present in the whole volume of the suspension. In contrast, in DIANA, a mixture of the compounds with the detection probe is added to the immobilized insulin receptor only present on the surface of the assay well. This may lead to a lower availability of binding sites for the compounds or the probe. Despite the non-covalent nature of the detection probe binding, this factor may play a role because insulin DIANA is sensitive to the order of added reaction components, unlike other setups. We tested both settings in the optimization phase of DIANA, i.e. adding ligands first to the insulin receptor, or the probe and compared the results to the common procedure we employ, that is, premixed ligands with the probe. In case ligands were applied first, we obtained lower K_d_ values for the inhibitors than with the mixture. In concert with that, the calculated K_d_ values were higher when the probe was added first than when the mixture was used. These effects most likely occur as a consequence of the unusual proportion of the detection probe. Insulin as a ligand for a detection probe is considerably larger than other ligands we commonly conjugate with the DNA oligonucleotide to synthesize a detection probe. Furthermore, the conformation of the insulin receptor might also play a role. While the radioligand cell-based assay employs the endogenously expressed receptor on the cell membrane, DIANA relies on the use of a recombinant receptor.

In addition, we identified several compounds as hits in manual DIANA post HTS that were not validated by the orthogonal assay. Hence, we carefully analyzed their structural motifs to understand the interference. The compounds contain aromatic cores and some of them also a positive charge, which leads to their binding to the DNA oligonucleotide of the detection probe and likely intercalation interfering with the subsequent qPCR. Some of the compounds were synthesized with the objective of labeling DNA, so this interference was expected. In our previous HTS experiments with other targets than the insulin receptor, such compounds were already identified as false-positive hits, but not to such an extent as in this assay (Simova et al.). Also in this case, we presume that this is due to the unique size of the detection probe. The two insulin molecules may certainly pose a steric hindrance, which then adds to the interfering effect of the compounds. It is also conceivable that the insulin molecules may be susceptible to the presence of certain compounds. Denaturation of one or both insulin moieties can negatively affect the qPCR leading to a false-positive result. With this detection probe, we aimed to push the boundaries of what we had commonly used for DIANA probe in the past. Therefore, we meticulously analyzed all hits with an orthogonal assay.

## Conclusion

Concerning sensitivity, both methods are comparable as can be demonstrated by the measurement of insulin K_d_ and K_d_ values of insulin analogs. IR binding assay uses radioactive materials, which require special handling, storage, and disposal procedures. As opposed to that, DIANA does not utilize radioactive substances, so it is easier and safer to handle. IR binding assay is a relatively time-consuming method due to the need for cell preparation over days. DIANA on the other hand is a fast assay that may be upscaled easily requiring the same amount of time, allowing screening of thousands of compounds within a few hours in the fully automated format. IR binding is affected by a larger measurement error due to pipetting by hand than DIANA performed by pipetting robots. The IR binding assay is the more costly method of the two as it utilizes radioactive material that itself is more expensive also concerning its disposal. DIANA in comparison is the more cost-effective alternative requiring only non-radioactive material. Only internal controls are necessary for interplate accuracy assessments. Moreover, HTS laboratories are often not authorized for work with radioactive substances, which poses a risk of contamination of the whole platform. Concerning quantification, DIANA is again the more reliable and straightforward method because IR binding assay relies on measuring radiation, which may pose issues with background noise and decay over time.

In summary, we propose DIANA as an excellent alternative to the radioactive IR binding assay with distinct merits over the latter method.

## Supporting information

Supplementary information updated

## Supporting information

The authors have cited additional references within the Supporting information.^[26]^

## Acknowledgements

The work of JS, KC, LZ, JJ and PS was supported by the program OP VVV funded by the Czech Ministry of Education, Youth and Sports. The work of JS and PS was supported by the Czech Science Foundation (ID Project No. 23-05642S). The work of JJ and LZ was supported by the project National Institute for Research of Metabolic and Cardiovascular Diseases (Program EXCELES, ID Project No. LX22NPO5104, Funded by the European Union – Next Generation EU), and by the Academy of Sciences of the Czech Republic (Research Project RVO:52, support to the Institute of Organic Chemistry and Biochemistry).

## Table of contents

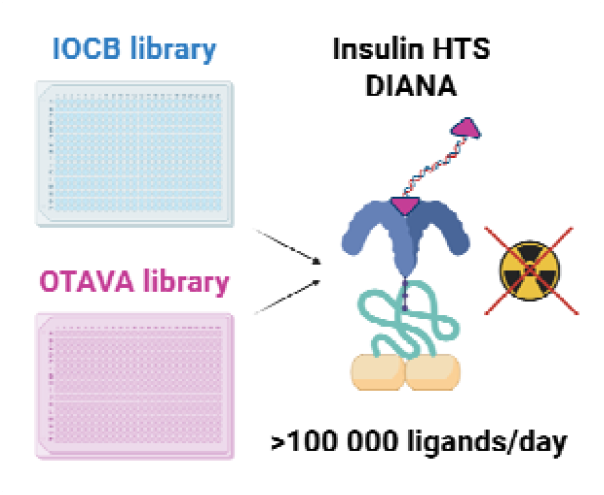

Created in BioRender. Stanurova, J. (2025) https://BioRender.com/qwsrljx

We present a non-radioactive high-throughput screening method for insulin receptor ligands. DIANA (DNA-linked inhibitor antibody assay) is faster, cheaper, easily scalable and most importantly safer than the commonly used radioligand receptor binding assay. In this work we screened two compound libraries confirming the sensitivity of the assay even in pooled format and verified our results with the radioligand receptor binding assay.

## Notes

### Competing Interest Statement

The authors have declared no competing interest.

### Summary of Updates

This version is a full update of the previous one. It contains updated results, compound synthesis and discussion. Supplementary information was added.

