## Supplementary information updated for "A Rapid and High-Sensitivity Non-radioactive Screening Assay for Insulin Receptor Ligands"

### Supporting information

#### 1. DIANA parameters

**A**

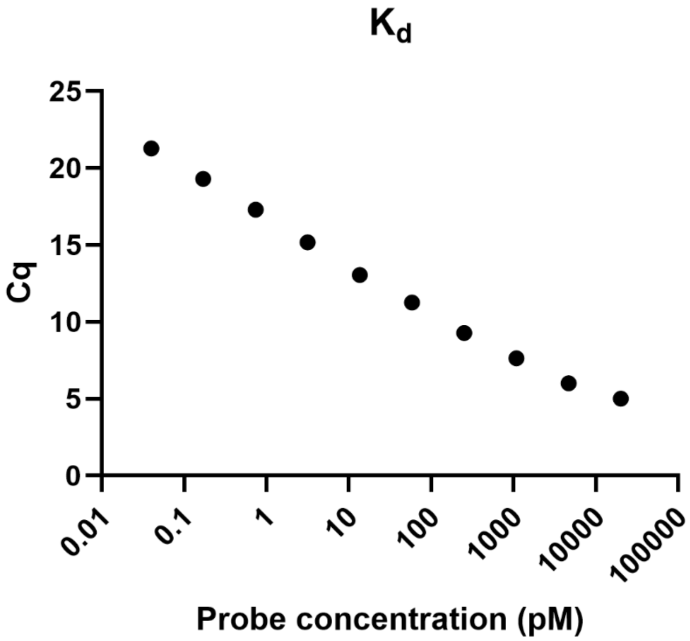

**B**

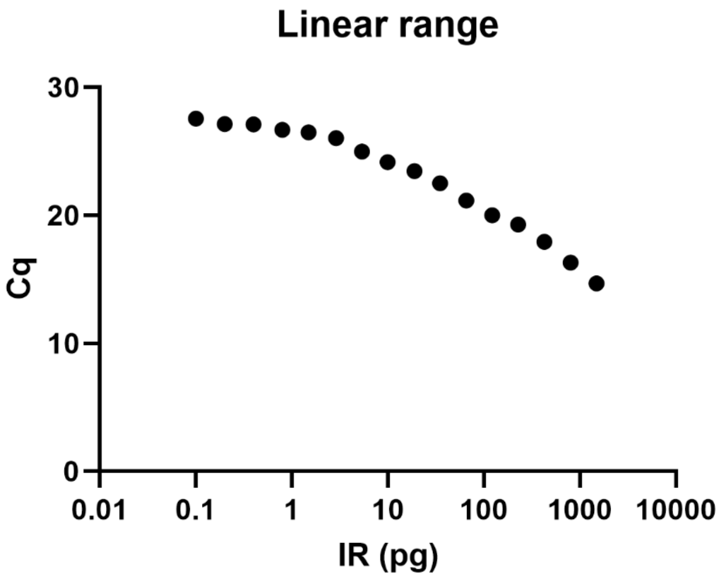

**C**

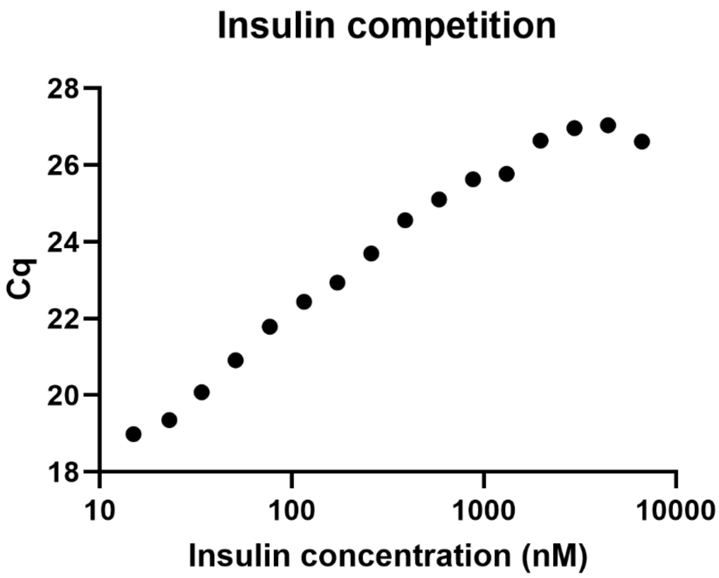

**Figure S1. A.** $K_d$ measurement. $K_d$ value of the bivalent probe was determined by DIANA. The probe was added to the insulin receptor in a dilution series (40 fM to 20 μM) and the amount of bound probe was monitored by RT-qPCR. The $K_d$ of the probe was calculated to be $K_d$ = 0.6 ± 0.1 nM. Cq = cycle of quantification. **B.** Linear range assessment. A serial dilution of the insulin receptor ectodomain ranging from 0.1 to 1500 pg was detected with 100 pM probe. Reliable detection of the insulin receptor ectodomain is possible even at 1.5 pg per well. **C.** Competition of the probe with a serial dilution of free insulin. Insulin mixed with the probe was added to the insulin receptor in concentrations of 15 to 15000 nM. The probe was successfully competed out, indicating non-covalent binding. The wide dynamic range of DIANA allows calculation of the insulin $K_d$ from Cq values across almost three orders of magnitude and thus insulin dilution is not required.

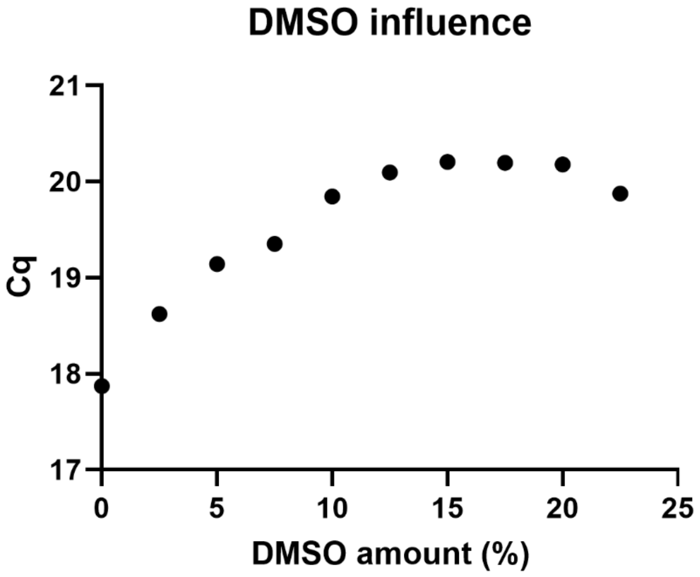

**Figure S2.** Determination of DMSO influence on the assay. DMSO was increased by 2.5 % in each step up to 22.5 %.

#### 2. Synthetic strategy for the preparation of insulin analogs LZ-113, LZ-120 and LZ-128

The first step involves preparing non-natural amino acid building blocks **S4**, **S5** and *ent*-**S5** (Scheme S1) for the solid-phase synthesis of peptide precursors and the subsequent enzymatic semisynthesis of insulin analogs. These blocks are designed either to replace the original amino acid at position B26 in full-length insulin analog **LZ-113** or to terminate the B26 site in *des*(B27–B30)-shortened analogs **LZ-120** and **LZ-128** with a C-terminal carboxamide functional group.

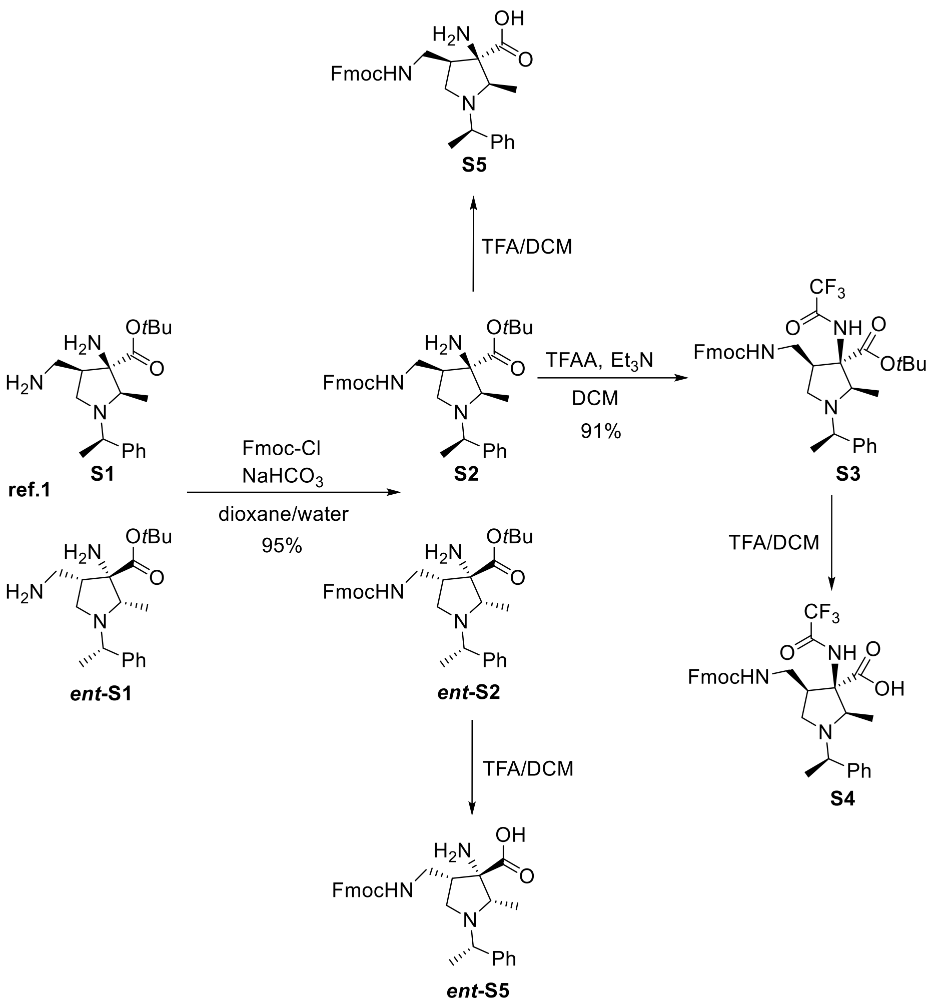

**Scheme S1.** Synthetic access to triamino acid building blocks **S4**, **S5** and *ent*-**S5**. The starting compounds **S1** and ***ent*-S1** were prepared as described in ref. 1.

The synthesis of building blocks begins with free diamine **S1** or its enantiomer ***ent*-S1**, which were prepared according to the literature.[1] The γ-amino group of these compounds was protected with Fmoc-Cl, yielding building blocks **S2** and ***ent*-S2**, respectively. The γ-Fmoc-protected amino acid **S2**, bearing a free α-amino group, was subsequently converted to trifluoroacetamide **S3** using TFAA. Finally, the *tert*-butyl groups of **S2**, *ent*-**S2** and **S3** were removed using TFA in DCM to yield **S5**, *ent*-**S5** and **S4**, respectively. These non-natural building blocks were used for the solid phase synthesis of precursor peptides for the enzymatic semisynthesis of insulin analogs. The solid-phase synthesis of peptides was performed as described in detail by Lubos et al.[2]

Block **S4** was subsequently incorporated into the octapeptide GFF-X-TP($K_{Pac}$)T at position X, replacing the original Tyr (corresponding to position B26 in insulin). The peptide was assembled on Wang resin preloaded with Fmoc-Thr(*O*tBu). $K_{Pac}$ refers to Fmoc-εN-phenylacetyl-Lys (ref. [3]). Next, the resulting peptide was attached *via* its N-terminal Gly residue to [*des*(B23–B30)]-insulin (**DOI**) by trypsincatalyzed semisynthesis[3] to generate an insulin analog precursor. Finally, the phenylacetyl group was cleaved with penicillin G acylase, as described by Žáková et al.[3], to yield analog **LZ-113**.

Blocks **S5** and ***ent*-S5** were incorporated at position X of the tetrapeptides GFFX using Rink Amide AM resin. Both **S5** and ***ent*-S5** contain a sterically hindered free amino group at α position. Interestingly, in both peptide products, this free α-amino group was unexpectedly acylated with glycine, likely during the coupling of the N-terminal amino acid (Figure S3). These peptides were subsequently used for the semisynthesis of insulin analogs **LZ-120** and **LZ-128**, following the same trypsin-mediated ligation described above.

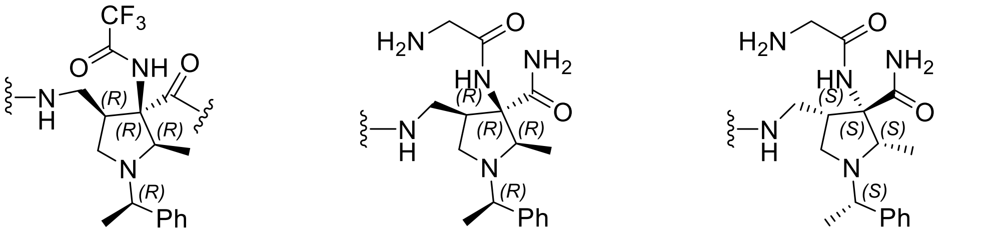

**Figure S3.** Unusual amino acids at the position B26 of insulin analogs **LZ-113**, **LZ-120** and **LZ-128**. For details, see below.

#### General experimental information

Reactions not involving aqueous conditions were performed in flame-dried glassware under an argon atmosphere. Solvents and additives were dried prior to use according to standard procedures. TLC analyses were performed on POLYGRAM SIL G/UV254 plates. Chromatographic separations were carried out on silica gel 60 (Fluka, 230-400 mesh) either manually or on a CombiFlash® NextGen 300+ instrument. ESI mass spectra were obtained on Thermo Fisher Scientific LCQ Fleet spectrometer, sample concentration approx. 1 µg/mL, spray voltage pos. mode: 3.3 kV. HRMS spectra were measured on Waters Q-Tof micro spectrometer, resolution: 100000. $^{1}H$ and $^{13}C$ NMR spectra were recorded on Bruker Avance III™ 400, 500 or 600 spectrometers operating at 400, 500 or 600 MHz for $^{1}H$ NMR and 100.1, 125.7 or 150.9 MHz for $^{13}C$ NMR. Temperature-dependent spectra were recorded on a Bruker Avance II™ 500 MHz instrument. Full assignment of $^{1}H$ and $^{13}C$ signals was achieved by a combination of 2D experiments ($^{1}H,^{1}H$-COSY; $^{1}H$-$^{13}C$ HMBC; $^{1}H$-$^{13}C$ HSQC). $CDCl_3$ was dried over 3Å molecular sieves prior to the measurements. IR spectra of the synthetized compounds were measured on Bruker AlPHA-FT-IR spectrometer (4 $cm^{-1}$ spectral resolution, Happ–Genzel apodization function, 64 scans) as neat samples using an ATR device equipped with diamond crystal in the 4000–600 $cm^{-1}$ spectral range.

#### 3. Experimental data and characterization

***tert*-Butyl (2*R*,3*R*,4*R*)-3-amino-4-(aminomethyl)-2-methyl-1-((*R*)-1-phenylethyl)pyrrolidine-3-carboxylate (S1)** and ***tert*-butyl (2*S*,3*S*,4*S*)-3-amino-4-(aminomethyl)-2-methyl-1-((*S*)-1-phenylethyl)pyrrolidine-3-carboxylate (*ent*-S1)**

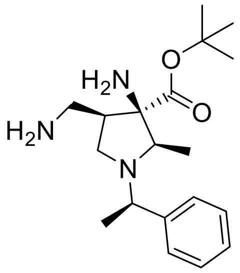

The compound was prepared in both enantiomeric forms (**S1** and *ent*-**S1**) according to the literature, the analytical data are in agreement.[1]

***tert*-Butyl (2*R*,3*R*,4*R*)-4-(((((9H-fluoren-9-yl)methoxy)carbonyl)amino)methyl)-3-amino-2-methyl-1-((*R*)-1-phenylethyl)pyrrolidine-3-carboxylate (S2)** and ***tert*-butyl (2*S*,3*S*,4*S*)-4-(((((9H-fluoren-9-yl)methoxy)carbonyl)amino)methyl)-3-amino-2-methyl-1-((*S*)-1-phenylethyl)pyrrolidine-3-carboxylate (*ent*-S2)**

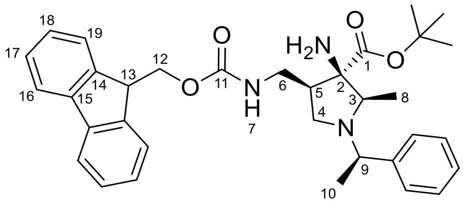

Fmoc-Cl (0.89 g, 3.44 mmol) in dioxane (10 mL) was dropwise added to a stirred solution of diamine **S1** (1.14 g, 3.44 mmol) and NaHCO3 (0.84 g, 6.88 mmol) in dioxane (30 mL) and water (10 mL) at 0 °C and the reaction mixture was stirred for 20 min. The reaction mixture was warmed to r.t., diluted with water (20 mL) and extracted with EtOAc (3x50 mL). The organic layer was dried over $Na_2SO_4$, filtered and the solvents were evaporated in vacuo. The residue was purified by flash column chromatography ($CHCl_3$/MeOH 50:1 to 10:1 gradient) to afford 1.82 g (95%) of protected amine **S2** as an off-white amorphous solid. $[\alpha]_D^{20}$: −25.5 (c 0.25, $CHCl_3$); IR ν[$cm^{-1}$]: 648, 741, 912, 1007, 1079, 1153, 1257, 1342, 1373, 1454, 1515, 1726, 2941, 2982, 3074, 3379; $^1H$ NMR (401 MHz, $C_6D_6$) δ 7.57 (dd, *J* = 7.2, 2.9 Hz, 2H, H-16), 7.51 (d, *J* = 7.3 Hz, 1H, H-19), 7.44 (d, *J* = 7.3 Hz, 1H, H-19), 7.36-7.32 (m, 1H, Ar), 7.26-7.08 (m, 8H, Ar, H-17, H-18), 5.39 (bs, 1H, H-7), 4.54 (dd, *J* = 10.6, 6.7 Hz, 1H, H-12a), 4.23 (dd, *J* = 10.6, 7.4 Hz, 1H, H-12b), 4.07 (t, *J* = 7.0 Hz, 1H, H-13), 3.71 (q, *J* = 6.7 Hz, 1H, H-9), 3.36 (ddd, *J* = 13.0, 7.6, 5.2 Hz, 1H, H-6a), 3.08-2.99 (m, 1H, H-6b), 3.03 (q, *J* = 6.2 Hz, 1H, H-3), 2.96-2.86 (m, 1H, H-5), 2.44 (t, *J* = 9.8 Hz, 1H, H-4a), 1.98 (dd, *J* = 9.6, 6.8 Hz, 1H, H-4b), 1.44 (s, 2H, $NH_2$), 1.32 (s, 9H, *t*Bu), 1.02 (d, *J* = 6.7 Hz, 3H, H-8), 0.76 (d, *J* = 6.2 Hz, 3H, H-10); $^{13}C$ NMR (101 MHz, $C_6D_6$) δ 173.5 (C, C-1), 156.3 (C, C-11), 144.9 (C, Ar), 144.6 (C, C-14), 141.8 (C, C-15),

128.4 (CH, Ar), 127.8 (CH, C-17), 127.3 (CH, C-18), 127.0 (CH, Ar), 125.8 (CH, Ar), 125.5 (CH, C-19), 120.2 (CH, C-16), 81.1 (C, *t*Bu), 67.8 (C, C-2), 66.7 ($CH_2$, C-12), 64.6 (CH, C-3), 55.2 (CH, C-9), 47.9 (CH, C-13), 47.2 ($CH_2$, C-4), 44.2 (CH, C-5), 41.2 ($CH_2$, C-6), 27.9 ($CH_3$, *t*Bu), 12.7 (CH, C-10), 11.3 (CH, C-8); MS (ESI+) m/z, (%): 556 (100, $[M+H]^+$), 578 (15, $[M+Na]^+$); HRMS (ESI+) m/z: $[M+H]^+$ Calcd for $C_{34}H_{42}N_3O_4$ 556.3170; Found 556.3170.

The other enantiomer *ent*-**S2** was prepared from free diamine *ent*-**S1** and the analytical data are in agreement.

***tert*-Butyl (2*R*,3*R*,4*R*)-4-(((((9H-fluoren-9-yl)methoxy)carbonyl)amino)methyl)-2-methyl-1-((*R*)-1-phenylethyl)-3-(2,2,2-trifluoroacetamido)pyrrolidine-3-carboxylate (S3)**

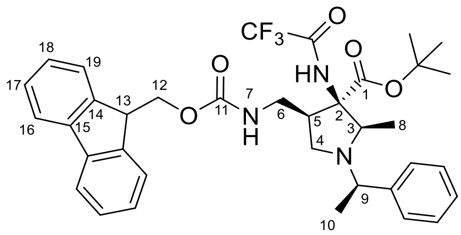

$Et_3N$ (171 µL, 1.23 mmol) was dropwise added to a stirred solution of amine **S2** (383 mg, 0.69 mmol) and TFAA (105 µL, 0.75 mmol) in DCM (1 mL) at 0 °C and the reaction mixture was warmed to r.t. and stirred overnight. The reaction mixture was diluted with water (5 mL) and extracted with EtOAc (3x15 mL). The organic layer was dried over $Na_2SO_4$, filtered and the solvents were evaporated *in vacuo*. The residue was purified by flash column chromatography (hexane/EtOAc + 1% TEA 5:1 to 2:1 gradient) to afford 409 mg (91%) of amine **S3** as an off-white amorphous solid. $[\alpha]_D^{20}$: −17.9 (c 0.2, $CHCl_3$); IR ν[$cm^{-1}$]: 621, 701, 741, 759, 844, 877, 1025, 1051, 1158, 1212, 1257, 1370, 1393, 1450, 1478, 1518, 1724, 2875, 2936, 2977, 3029, 3065, 3405; $^1$H NMR (400 MHz, $CDCl_3$) δ 7.78-7.73 (m, 2H, H-16), 7.65-7.60 (m, 2H, H-19), 7.42-7.24 (m, 9H, H-17, H-18, Ar), 6.77 (s, 1H, $NHCOCF_3$), 5.83 (dd, *J* = 8.9, 3.6 Hz, 1H, H-7), 4.38 (dd, *J* = 10.5, 7.5 Hz, 1H, H-12a), 4.32 (dd, *J* = 10.5, 7.0 Hz, 1H, H-12b), 4.21 (t, *J* = 7.2 Hz, 1H, H-13), 4.09 (q, *J* = 6.6 Hz, 1H, H-9), 3.43 (ddd, *J* = 14.2, 9.1, 4.0 Hz, 1H, H-6a), 3.29-3.19 (m, 2H, H-3, H-5), 2.92 (t, *J* = 10.0 Hz, 1H, H-4a), 2.57 (ddd, *J* = 14.3, 10.2, 3.7 Hz, 1H, H-6b), 1.99 (dd, *J* = 9.9, 8.2 Hz, 1H, H-4b), 1.48 (s, 9H, *t*Bu), 1.33 (d, *J* = 6.7 Hz, 3H, H-10), 1.15 (d, *J* = 6.3 Hz, 3H, H-8); $^{13}$C NMR (101 MHz, $CDCl_3$) δ 166.7 (C, C-1), 157.8 (q, *J* = 37.0 Hz, C, $COCF_3$), 156.6 (C, C-11), 144.3 (C, C-14), 142.6 (C, Ar), 141.4 (C, C-15), 128.6 (CH, Ar), 127.7 (CH, C-17), 127.4 (CH, Ar), 127.3 (CH, Ar), 127.2 (CH, H-18), 125.4 (CH, H-19), 120.0 (CH, H-16), 116.0 (q, *J* = 288.3 Hz, C, $CF_3$), 83.2 (C, *t*Bu), 68.7 (C, C-2), 66.9 ($CH_2$, C-12), 63.2 (CH, C-3), 54.1 (CH, C-9), 47.4 (CH, C-13), 45.7 ($CH_2$, C-4), 43.1 (CH, C-5), 40.1 ($CH_2$, C-6), 27.8 ($CH_3$, *t*Bu), 12.8 ($CH_3$, C-8), 10.1 ($CH_3$, C-10); $^{19}$F NMR (377 MHz, $CDCl_3$) δ —75.67; MS (ESI+) m/z, (%): 596 (100, $[M+H-C_4H_8]^+$), 652 (81, $[M+H]^+$); HRMS (ESI+) m/z: $[M+H]^+$ Calcd for $C_{36}H_{41}N_3O_5F_3$ 652.2993; Found 652.2992.

#### 5. NMR spectra

***tert*-Butyl (2*R*,3*R*,4*R*)-4-(((((9H-fluoren-9-yl)methoxy)carbonyl)amino)methyl)-3-amino-2-methyl-1-((*R*)-1-phenylethyl)pyrrolidine-3-carboxylate (S2)**

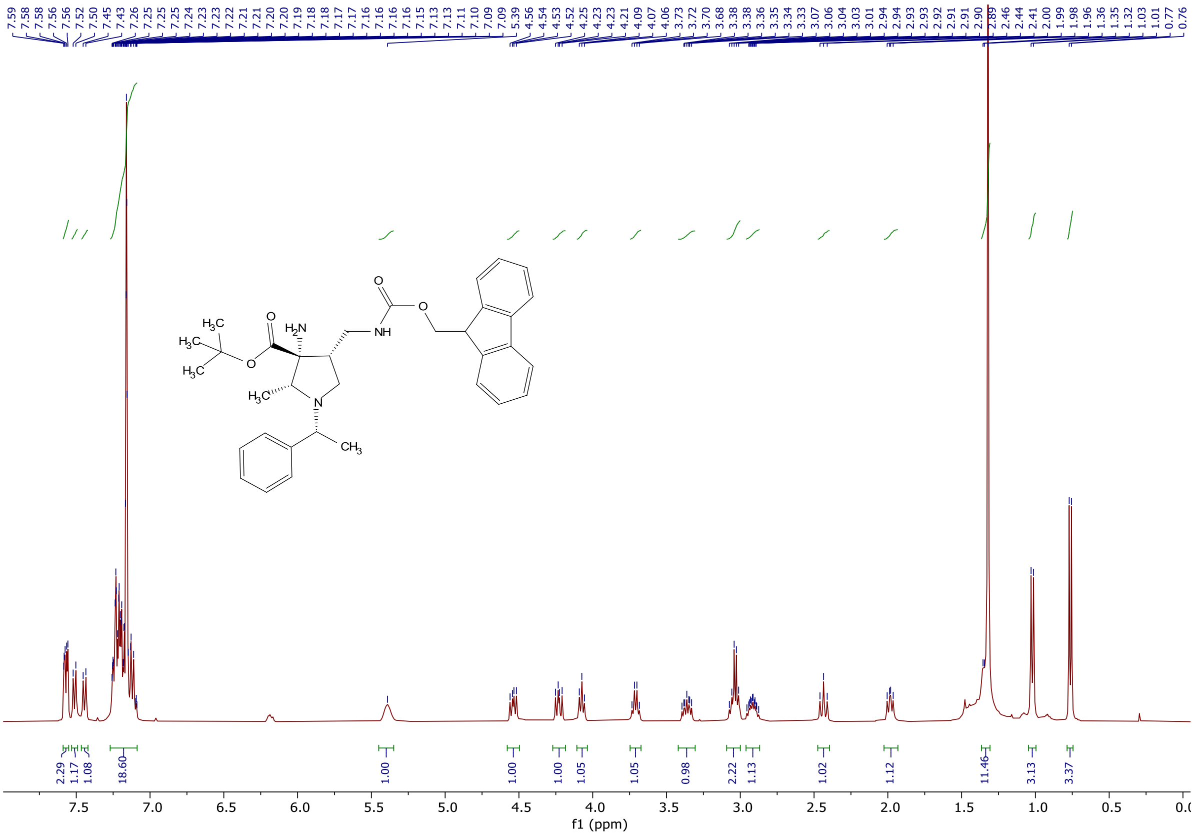

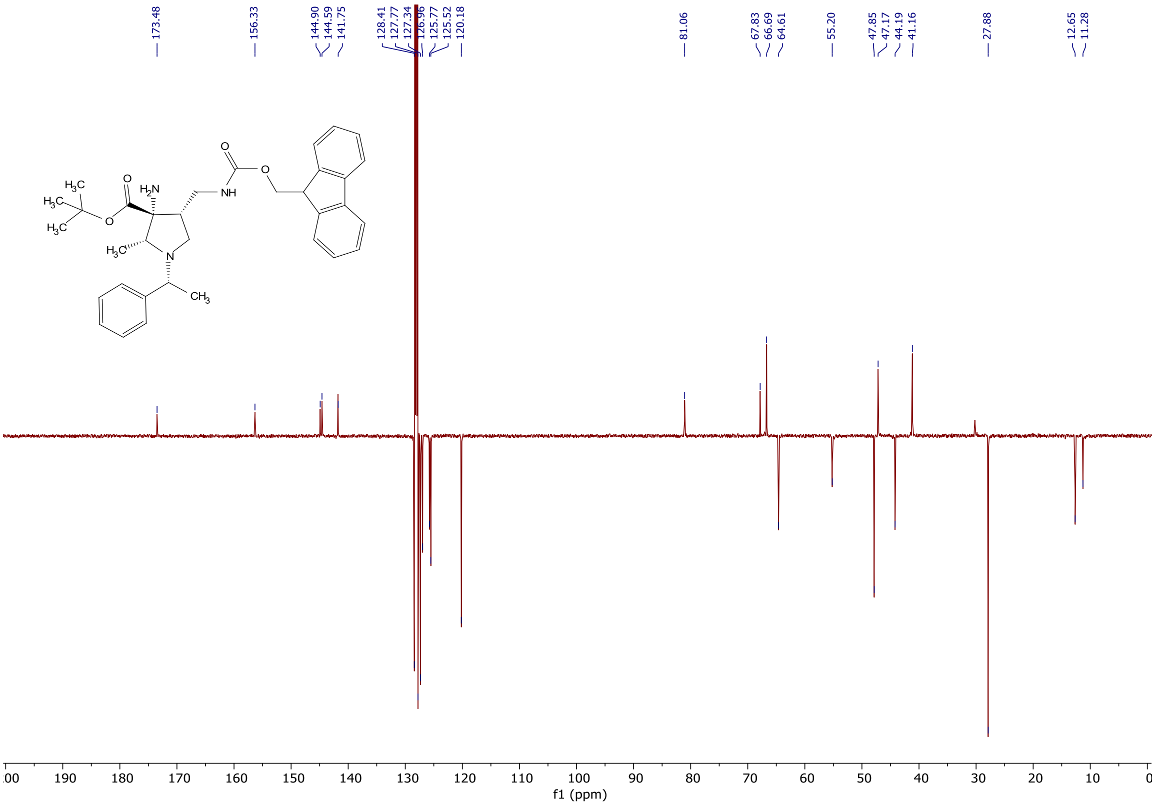

***tert*-Butyl (2*R*,3*R*,4*R*)-4-(((((9H-fluoren-9-yl)methoxy)carbonyl)amino)methyl)-2-methyl-1-((*R*)-1-phenylethyl)-3-(2,2,2-trifluoroacetamido)pyrrolidine-3-carboxylate (S3)**

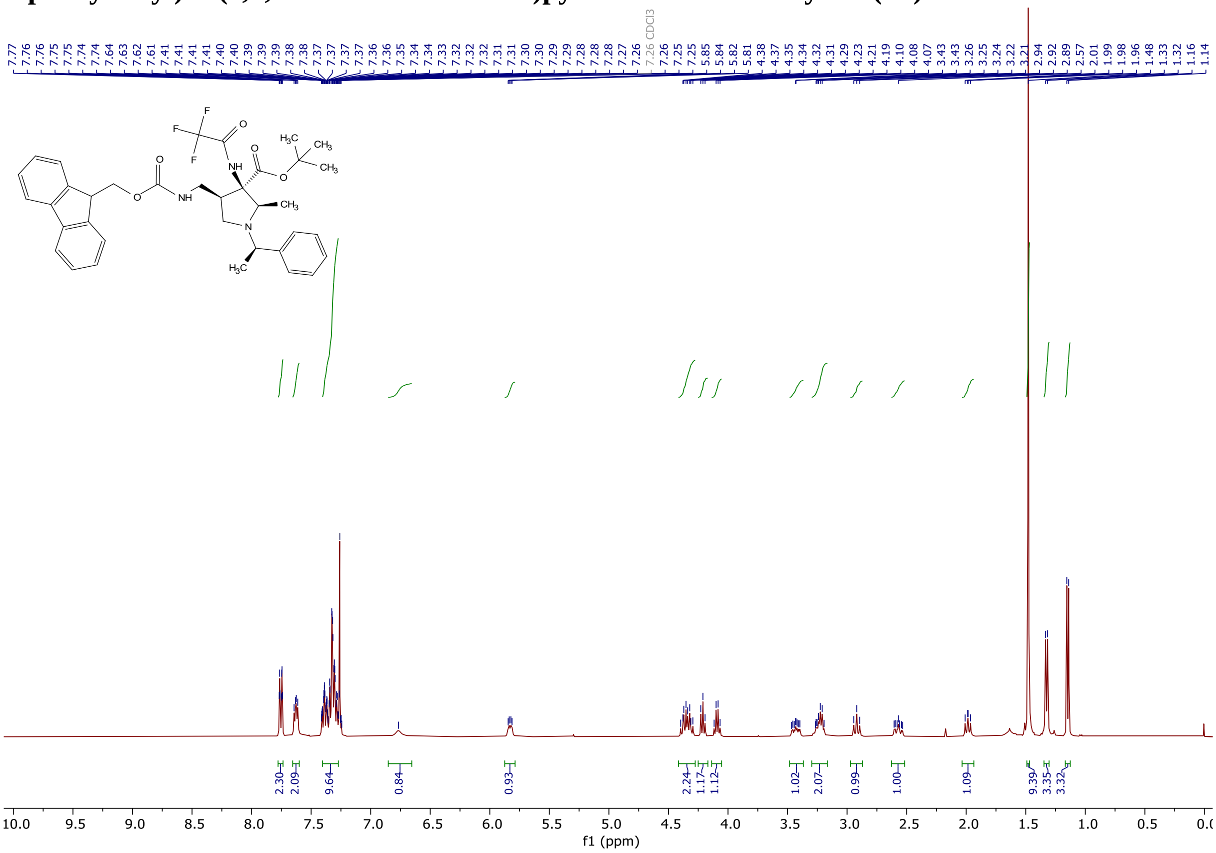

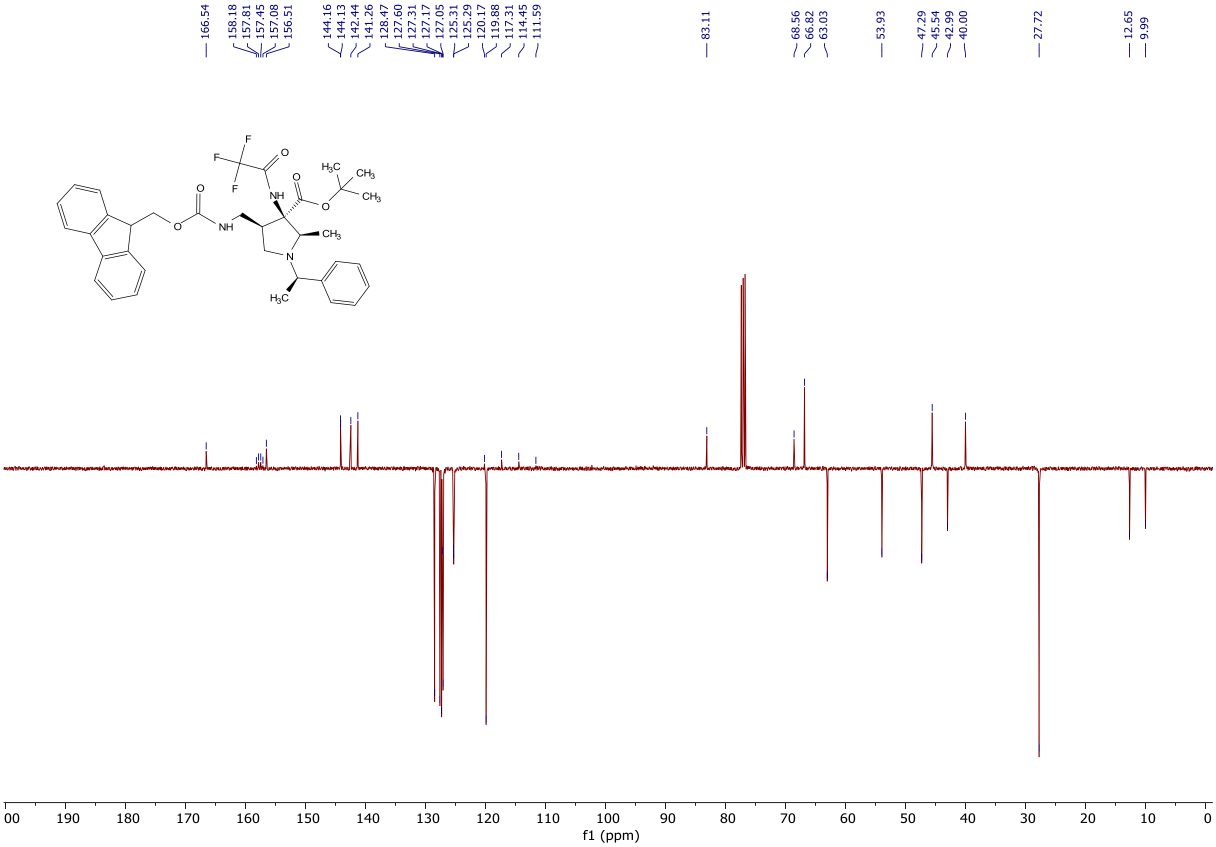

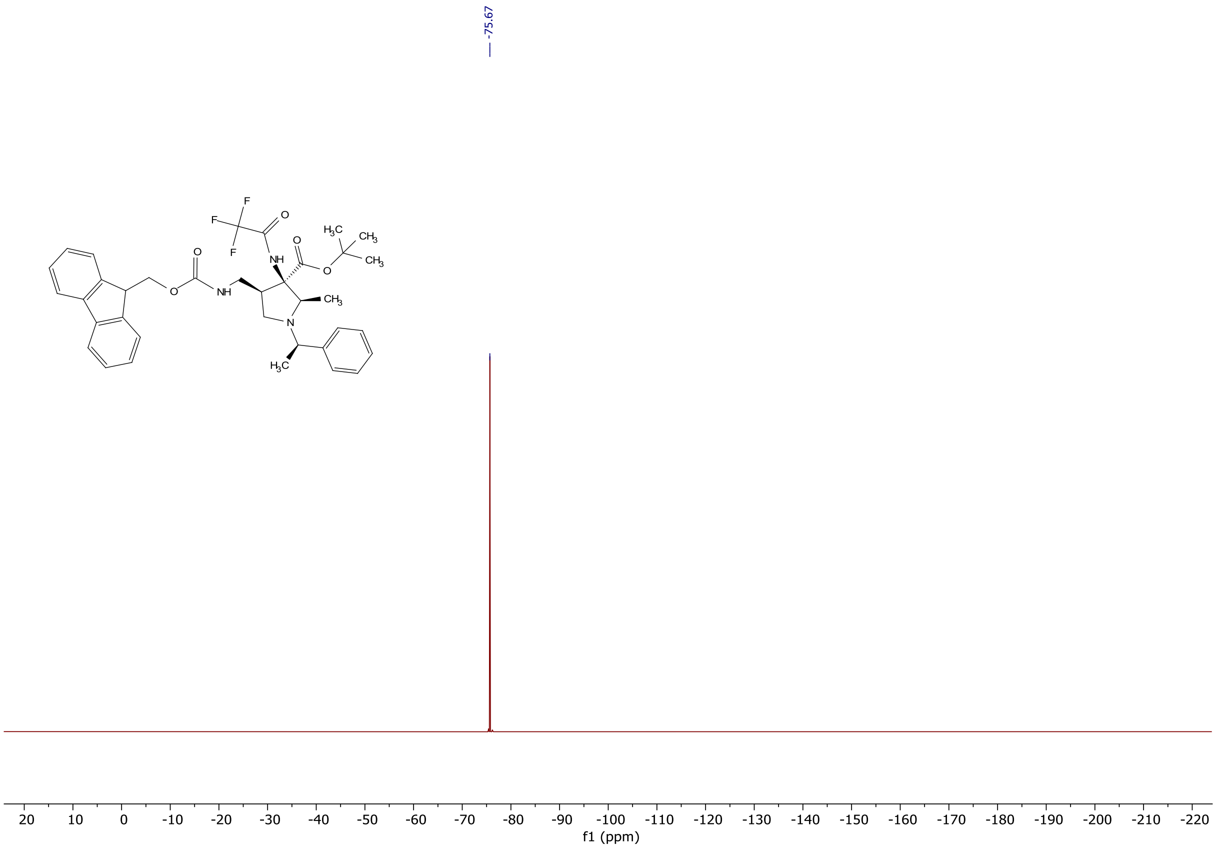
-75.67
F
F
F
O
NH
O
H3C
CH3
CH3
O
O
O
NH
CH3
N
H3C
20 10 0 -10 -20 -30 -40 -50 -60 -70 -80 -90 -100 -110 -120 -130 -140 -150 -160 -170 -180 -190 -200 -210 -220
f1 (ppm)
